# Development of novel apoptosis-assisted lung tissue decellularization methods

**DOI:** 10.1101/2020.05.21.109173

**Authors:** Young Hye Song, Mark A Maynes, Nora Hlavac, Daniel Visosevic, Kaitlyn O Daramola, Stacy L Porvasnik, Christine E Schmidt

## Abstract

Decellularized tissues hold great potential for both regenerative medicine and disease modeling applications. The acellular extracellular matrix (ECM)-enriched scaffolds can be recellularized with patient-derived cells prior to transplantation, or digested to create thermally-gelling hydrogels for 3D cell culture. Current methods of decellularization clear cellular components using detergents, which can result in loss of ECM proteins and tissue architectural integrity. Recently, an alternative approach utilizing apoptosis to decellularize excised murine sciatic nerves resulted in superior ECM preservation, cell removal, and immune tolerance *in vivo*. However, this apoptosis-assisted decellularization approach has not been optimized for other tissues with a more complex geometry, such as lungs. To this end, we developed an apoptosis-assisted lung tissue decellularization method using a combination of camptothecin and sulfobetaine-10 (SB-10) to induce apoptosis and facilitate gentle and effective removal of cell debris, respectively. Importantly, combination of the two agents resulted in superior cell removal and ECM preservation compared to either of the treatments alone, presumably because of pulmonary surfactants. In addition, our method was superior in cell removal compared to a previously established detergent-based decellularization protocol. Furthermore, thermally-gelling lung ECM proteins supported high viability of rat lung epithelial cells for up to 2 weeks in culture. This work demonstrates that apoptosis-based lung tissue decellularization is a superior technique that warrants further utilization for both regenerative medicine and disease modeling applications.

## 1. Introduction

Advances in tissue engineering have led to broad use of naturally-derived biomaterials such as collagen, fibrin, hyaluronan, and decellularized tissue extracellular matrix (ECM) scaffolds, for regenerative medicine as well as for ex vivo disease modeling^1–4^. Naturally-derived biomaterials offer unique advantages in both applications because they provide natural cell attachment sites and are biocompatible, biodegradable, and cause little to no immune response post-implantation^2^. Additionally, natural biomaterial scaffolds allow assessment of cell behavior in a physiologically relevant manner as two-dimensional (2D) substrate coatings and as three-dimensional (3D) cell culture scaffolds^1^. Studies have collectively demonstrated the importance of ECM-derived cues in cell response in healthy and diseased states, further highlighting the importance of environmental relevance when understanding cell behavior and translating the results to clinical applications.

Among different naturally-derived biomaterials, decellularized tissues and organs offer unique advantages in generating acellular scaffolds with high retention of the majority of tissue-specific ECM components while removing immunogenic cellular debris^5^. As such, decellularized tissues have been widely used to repair damaged tissues and replace lost or non-functional tissues/organs. Furthermore, recent studies have emerged in which the decellularized tissues are further processed to create 3D cell culture scaffolds for injectable delivery vehicles of cells, drugs, and pro-regenerative cytokines.^6–11^

This increased use of decellularized tissues necessitates a thorough review of current practices in tissue decellularization. The majority of current decellularization methods heavily relies on detergents to remove cellular debrisy^5^. Chemical processing often begins with water washes to induce necrosis of tissue-resident cells. Although chemical processing has achieved numerous pre-clinical and clinical successes, there still exists room for improvement. Necrosis results in rupture of cells and release of intracellular components that can cause immune response. To remove these immunogenic cellular debris, harsh washing steps are required, which further creates a problem of compromising tissue integrity^12^. Recently, an alternate approach was suggested in which tissues are decellularized by means of programmed cell death, or apoptosis, in which cells fragment into apoptotic bodies containing intracellular debris; these apoptotic bodies allow easier removal of cellular debris^13–15^. This approach does not rely on heavy use of detergents and water washes to induce cell death, but instead generates apoptotic bodies that are easier to remove via gentle washes. Recent publications on apoptosis-assisted decellularization of rat sciatic nerves^14^ and 3D *in vitro* tissue constructs^15^ have demonstrated excellent removal of cellular components and preservation of ECM proteins. Furthermore, *in vivo* implantation has shown immunocompetence of these scaffolds^14^. However, this approach has only been tested in a limited number of systems, including excised rat sciatic nerve and engineered 3D *in vitro* tissue cultures. In addition, the use of apoptosis-facilitated acellular tissue scaffolds as *in vitro* cell culture scaffolds has not yet been reported.

Therefore, the goal of this study was to expand the applicability of apoptosis-assisted tissue decellularization to a tissue type where complex micro- and macro-structure complicates efficient removal of cellular debris. To this end, we decellularized rat lung tissues using the apoptosis-inducing agent camptothecin, which has demonstrated success in decellularizing sciatic nerves via apoptosis^14^. We confirmed camptothecin-induced apoptosis via detection of DNA fragments. Our method results in great preservation of tissue ECM components (i.e., collagen types I and IV, laminin), while removing various intracellular components, including nuclear, cytoskeletal, and cell surface junction proteins. We further show that our method results in better overall removal of cellular components than a previously published detergents-based method^16^. Interestingly, a combination of camptothecin and sulfobetaine-10 (SB-10) zwitterionic detergent wash proved to be more effective than either reagents alone, presumably because of the pulmonary surfactants. Finally, we demonstrated that these acellular scaffolds can be digested to create 3D cell culture hydrogels, which exhibits thermally-gelling ECM hydrogel behavior and maintains high viability of encapsulated cells over a two-week time period. This study presents a novel method of generating acellular tissue scaffolds for 3D culture of cells in a native-like manner. Furthermore, this approach can be scaled up to generate acellular organs for transplantation or be used to improve injectable hydrogel systems *in vivo*.

## 2. Materials and Methods

### 2.1. Lung Procurement

Rat lung harvest procedure was approved by the Institutional Animal Care and Use Committee (IACUC) at the University of Florida. Lungs were procured as previously described^17^. Briefly, male and female Sprague-Dawley rats (250-300g, Charles River) were anesthetized with isoflurane vapor and systemically heparinized. Afterwards, the rib cages were opened to expose the heart and lungs. Lungs were blanched with ice-cold, sterile phosphate buffered saline (PBS) via intracardiac perfusion through the right ventricle. Individual lung lobes were placed in ice-cold, phenol red-free, sterile Dulbecco’s Modified Eagle Medium with F-12 nutrient mix (DMEM/F12, Gibco) and immediately transferred to ice-cold, sterile plastic petri dish. Sterile biopsy punches 4 mm in diameter were used to generate lung pieces. Lung pieces were then subjected to the following treatments: 1) fixation in 4% paraformaldehyde for immunohistochemical analysis of fresh lung pieces, 2) flash freezing in liquid nitrogen and lyophilization for biomolecular assays, and/or 3) immediate placement in camptothecin-containing medium to begin apoptosis-assisted decellularization.

### 2.2. Apoptosis-assisted Lung Decellularization

Decellularization of lung pieces via apoptosis is summarized in Figure 1. Briefly, lung pieces were placed in 20 µM camptothecin (Sigma-Aldrich) in DMEM/F12 with 1% penicillin/streptomycin (Gibco) and incubated for 48 hours at 37°C under agitation. Afterwards, lungs were washed in sterile hypertonic (4X) PBS at room temperature for 12 hours to a total of 4 washes. Next, the lungs were rinsed in 125 mM sulfobetaine-10 (SB-10, Sigma-Aldrich) for 7 hours, isotonic (1X) PBS for 1 hour twice, and statically incubated with 75 U/mL deoxyribonuclease (DNase) for 24 hours. Afterwards, lung pieces were rinsed in 1X PBS for 30 minutes 3 times, and purified water (Barnstead GenPure Water Purification Systems, Thermo Fisher) for 15 minutes twice. Next, the lung pieces were either fixed in 4% paraformaldehyde (Electron Microscopy Sciences) in PBS for 24 hours at 4°C for subsequent tissue processing, paraffin embedding, sectioning, and immunohistochemical analysis. Alternatively, they were flash frozen in liquid nitrogen and lyophilized for molecular assays and hydrogel formation. With the exception of the DNase step, lung pieces were agitated during incubation for each step using a vertical rotator.

**Figure 1.**
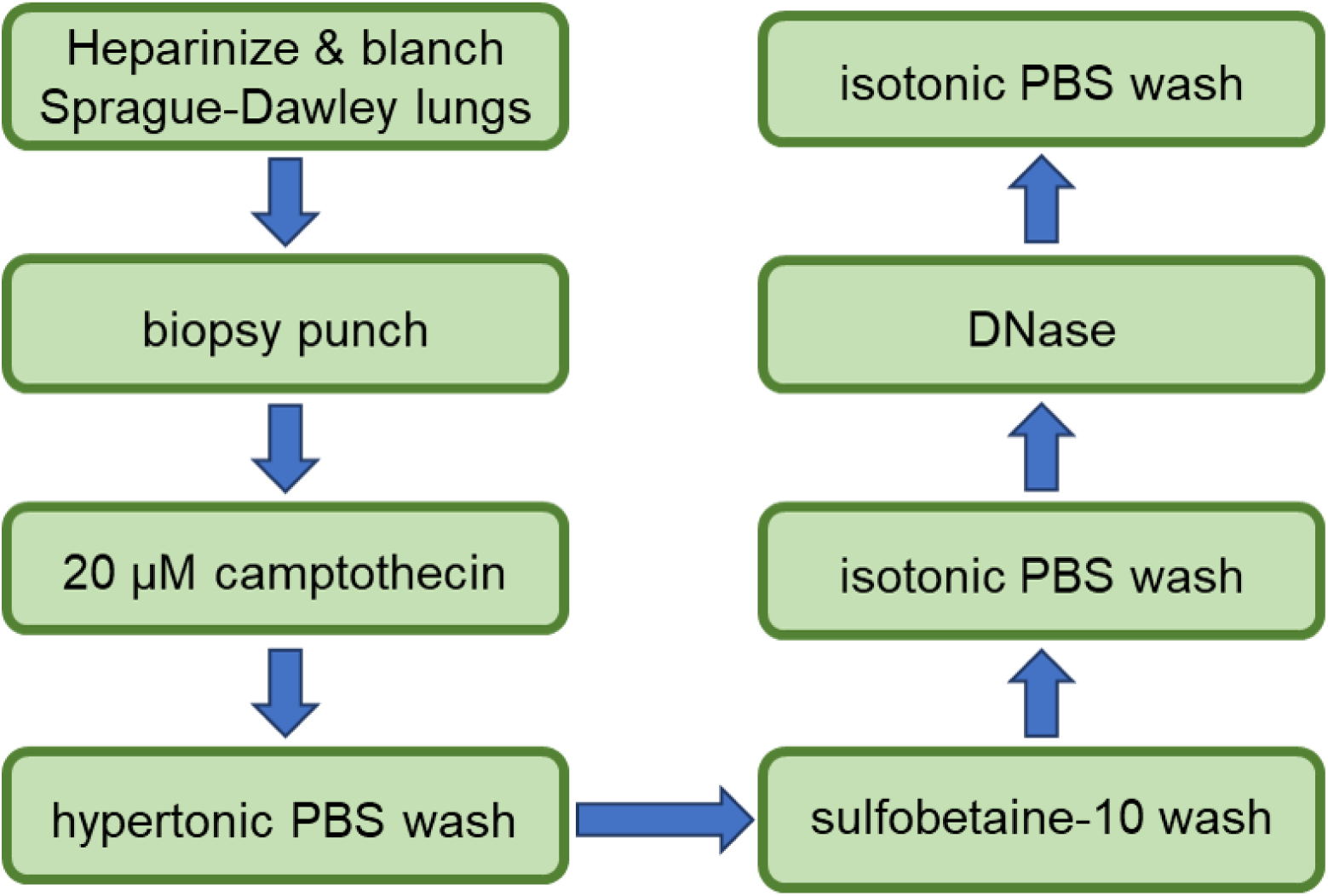
Overview of apoptosis-assisted lung tissue decellularization process.

To ensure that both camptothecin and SB-10 are needed for successful decellularization, two variations of the aforementioned method were carried out. First, in the “Campto Only” method, all the decellularization steps minus the SB-10 wash were done. In the “SB-10 Only” method, all the steps except the initial camptothecin treatment were performed.

### 2.3. Detergent-based Lung Decellularization

To compare the efficacy of the apoptosis decellularization method, a previously published detergent-based decellularization method was used^16^. Briefly, lung pieces were placed in 50 mL conical tubes and washed with deionized water for 1 hour, 0.1% (v/v) Triton X-100 for 52 hours, water for another hour, 2% (w/v) sodium deoxycholate for 26 hours, water for 1 hour, DNase for 1 hour, water for 1 hour, 1M sodium chloride for 1 hour, water for 1 hour and then PBS for 1 hour and 20 minutes. Detergents washes were carried out at 4°C, and all other washes were carried out at room temperature^16^. All steps except the DNase treatment were performed with agitation.

### 2.4. Immunohistochemistry

After fixation, lung pieces were placed in tissue cassettes available in the University of Florida Molecular Pathology Core (MPC), stored in 70% ethanol at 4°C and submitted to the MPC for paraffin embedding and sectioning 5 µm in thickness. These slides were utilized for different staining. For all immunohistochemistry (IHC) procedures, the slides were deparaffinized by first incubating at 60°C oven for 30 minutes, and then subjected to a series of xylene and decreasing ethanol concentration washes.

For histological analysis, hematoxylin and eosin (H&E) staining was performed using Harris hematoxylin solution (Sigma-Aldrich) and eosin solution (Electron Microscopy Sciences). Apoptosis was assessed by DNA fragmentation staining using a commercially available kit (In Situ Cell Death Detection Kit, TMR red, Sigma-Aldrich).

For immunohistochemical analysis, rehydrated slides were subject to antigen retrieval using citric acid buffer at 120°C and 20 psi in a pressure cooker (DAKO). Afterwards, the slides were briefly rinsed in Tris buffered saline (TBS) and treated with a blocking buffer containing 3% (v/v) goat serum and 0.3% (v/v) Triton X-100 in TBS. The slides were then incubated with primary antibodies [CD31 (abcam), α-smooth muscle actin (α-SMA, abcam), collagen I (abcam), collagen IV (Sigma-Aldrich), and laminin (Sigma-Aldrich)] diluted in the blocking buffer for 24 hours at 4°C. Subsequently, the slides were rinsed in TBS, and incubated with secondary antibodies diluted in blocking buffer for 24 hours at 4°C. The next day, the slides were rinsed in TBS, incubated with 4′,6-diamidino-2-phenylindole (DAPI, Thermo Fisher), rinsed with TBS, and mounted using Fluoromount G (SouthernBiotech) and glass coverslips. After overnight drying, the coverslips were affixed using nail polish and the slides were imaged using Zeiss Axio Imager upright epifluorescence microscope.

Image analysis of IHC images were done using ImageJ from the National Institute of Health (Bethesda, Maryland). Each histological image was split into three color channels and the threshold of each channel was found. The total positive pixel area and mean gray value were determined for intracellular and extracellular components, respectively.

### 2.5. Biomolecular Content Quantification

Quantification of DNA, GAG, and collagen was performed on lyophilized lung tissues that underwent different decellularization methods as described above. DNA was extracted using a commercially available kit (Qiagen DNeasy Blood & Tissue Kit). Afterwards, double-stranded DNA was quantified using QuantiFluor dsDNA System (Promega). Glycosaminoglycan (GAG) content was measured using a Blyscan GAG assay kit (Biocolor UK). A hydroxyproline assay kit (Abcam) was used for collagen content measurement. For dry tissue weight measurements, lyophilized lung pieces were weighed.

### 2.6. Metabolic Activity Assessment

To assess metabolic activity of any residual lung-resident cells after decellularization, the lung pieces were collected right after harvest and also at the end of decellularization. The pieces were placed in DMEM/F12 supplemented with 10% (v/v) alamarBlue reagent (Thermo Fisher) and incubated at 37°C for 3 hours. The media were collected, and fluorescence was read with excitation at 540 nm and emission at 590 nm. Changes in fluorescence were used to correlate cell metabolic activity.

### 2.7. Lung ECM Hydrogel Formation

Lyophilized, decellularized lung pieces were placed in sterile scintillation vials and minced with a pair of sterile microscissors (Eriem Surgical). Lung pieces were digested with 1 mg/mL pepsin in 0.01M hydrochloric acid (both from Sigma-Aldrich) at a tissue concentration of 10 mg/mL. Digestion was carried out at room temperature on a stir plate with a small magnetic stir bar. The pre-gel solution was transferred to the autoclaved 2 mL microcentrifuge tube using sterile micropipette tips, pH was increased to 7.4 using sterile 1M NaOH and 1M HCl, and 10X PBS solution was added at 10% (v/v) final concentration. Afterwards, the pre-gel solution was pipetted into a circular silicone mold 4 mm in diameter and 500 µm in depth, and incubated at 37°C until gelation.

### 2.8. Rheology on Lung ECM Hydrogels

Rheological measurements of the apoptosis-mediated lung ECM hydrogels were characterized using an MCR 302 rheometer (Anton Paar) to determine viscoelastic properties. Each pre-gel sample was prepared using the same method described above; however, 150 µL pre-gel solution was pipetted into a silicone mold (4mm diameter, 2mm depth, Grace Bio-Labs) and incubated for 45 minutes before rheological measurements. The hydrogels were placed on a preheated sand blasted bottom plate at 37°C with a top plate lowered to a height of 1-1.8 mm such that it contacts the hydrogel surface. The hydrogel was placed on the bottom plate with purified water surrounding the entire ring edge to prevent hydrogel dehydration. An amplitude sweep was conducted at a constant frequency of 6.3 rad/s from 0.01 to 100% strain to determine the linear viscoelastic region (LVR) where the storage and loss moduli are constant. A strain value was chosen to conduct frequency sweep measurements on lung ECM hydrogels as this value was within the LVR. The LVR was between 0.01% and 0.158% strain as determined from the amplitude sweeps; therefore, we chose a 0.1% strain to conduct the frequency sweeps.

### 2.9. 3D Culture of Lung Epithelial Cells in Lung ECM Hydrogels

CCL-149 rat lung epithelial cells (ATCC) were cultured in DMEM/F12 with 10% fetal bovine serum (FBS, R&D Systems) and 1% penicillin/streptomycin (P/S, Gibco) in a 37°C incubator with 5% CO_2_. When 85-90% confluent, cells were passaged and embedded in the lung ECM pre-gel solution at 2 million cells/mL. After gelation, the gels were individually placed in a 48-well plate and cultured in the CCL-149 growth media. At days 1, 4, 7 and 14, a live/dead assay was performed on CCL-149 in lung ECM hydrogels using a commercially available kit (Thermo Fisher). A Zeiss confocal microscope L710 was used to visualize cells in lung ECM hydrogels. The ratio between live and dead cells was analyzed using ImageJ.

### 2.10. Statistical Analysis

GraphPad Prism was used to plot data and analyze statistical significance. Student’s t-test (for alamarBlue assay) and one-way analysis of variance (ANOVA) with decellularization methods as a factor were performed. Tukey’s post-hoc analysis was used to assess specific differences across all groups. In the case of non-parametric data (DAPI staining), a Wilcoxon method was used to compare each pair. All data are presented as mean ± standard deviation.

## 3. Results and Discussion

### 3.1. Histological Assessments Show Nuclear Removal, Tissue Preservation, and DNA Fragmentation in Lung Pieces

To validate successful decellularization of lung pieces via apoptosis, histological analysis of the lung pieces was performed at different stages of the decellularization process. To this end, lung pieces were fixed at different steps (“**Fresh”** immediately after harvest, “**Campto”** immediately after the 48-hour camptothecin treatment, “**SB-10”** immediately after the SB-10 wash, and “**End”** indicating the end of decellularization), paraffin embedded and sectioned, and stained using H&E. Furthermore, to ensure that the cells were undergoing apoptosis during camptothecin treatment, Fresh and Campto lung piece sections were stained for DNA fragments. H&E images show that the lung pieces were devoid of cell nuclei at the end of decellularization, indicating successful cell removal (Figure 2). Furthermore, the tissues maintained their architecture, as demonstrated in Figure 2 with the distinct alveolar structure preserved at all stages of lung decellularization. As for confirmation of apoptosis, lung-resident cells after camptothecin treatment were positive for DNA fragments, as confirmed with immunofluorescence images and quantification of DNA fragment-positive nuclei (Figure 3a-b).

**Figure 2.**
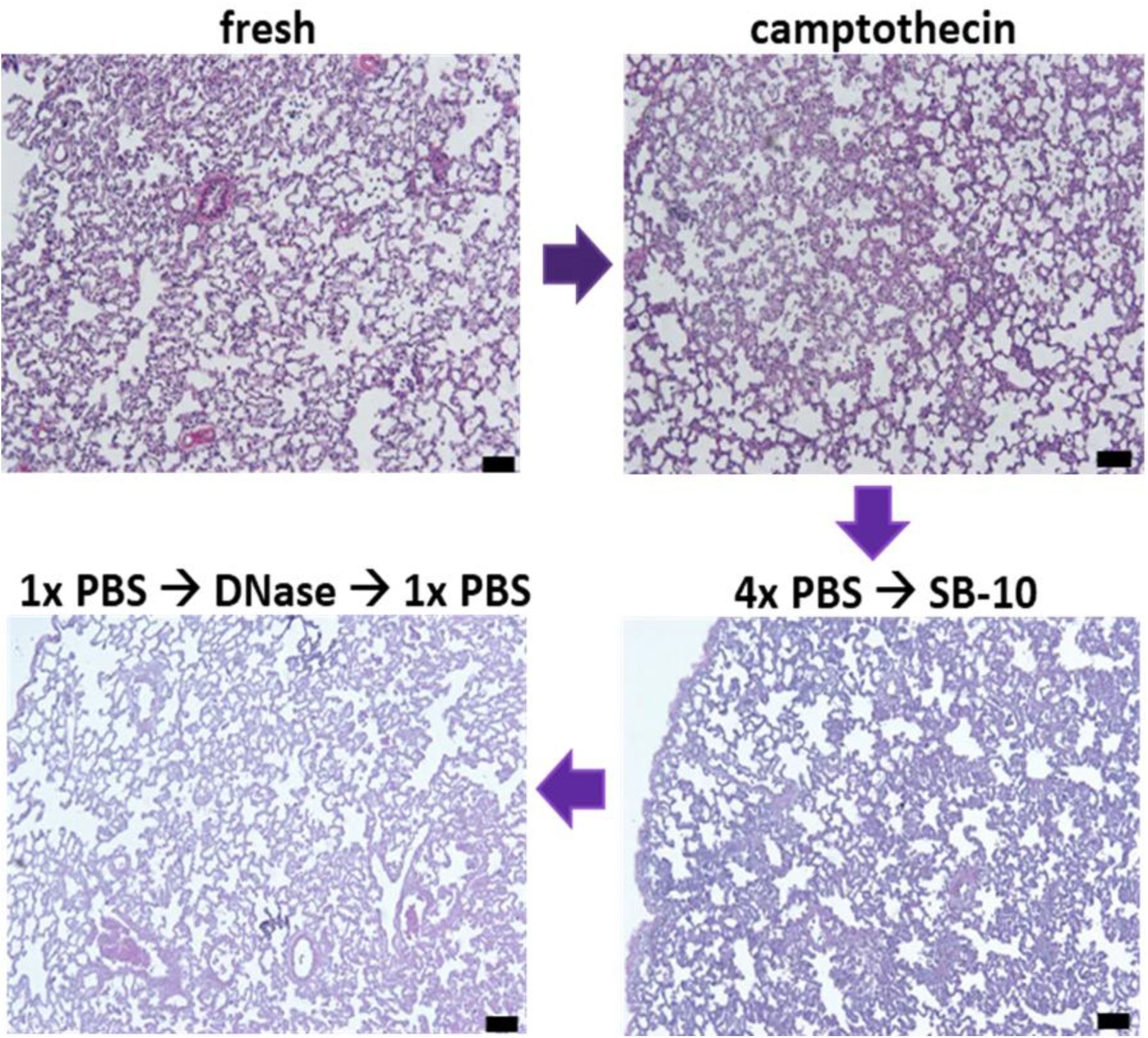
Hematoxylin and eosin (H&E) staining of paraffin-embedded lung tissue sections at different stages of the decellularization. Scale bar: 100 µm.

**Figure 3.**
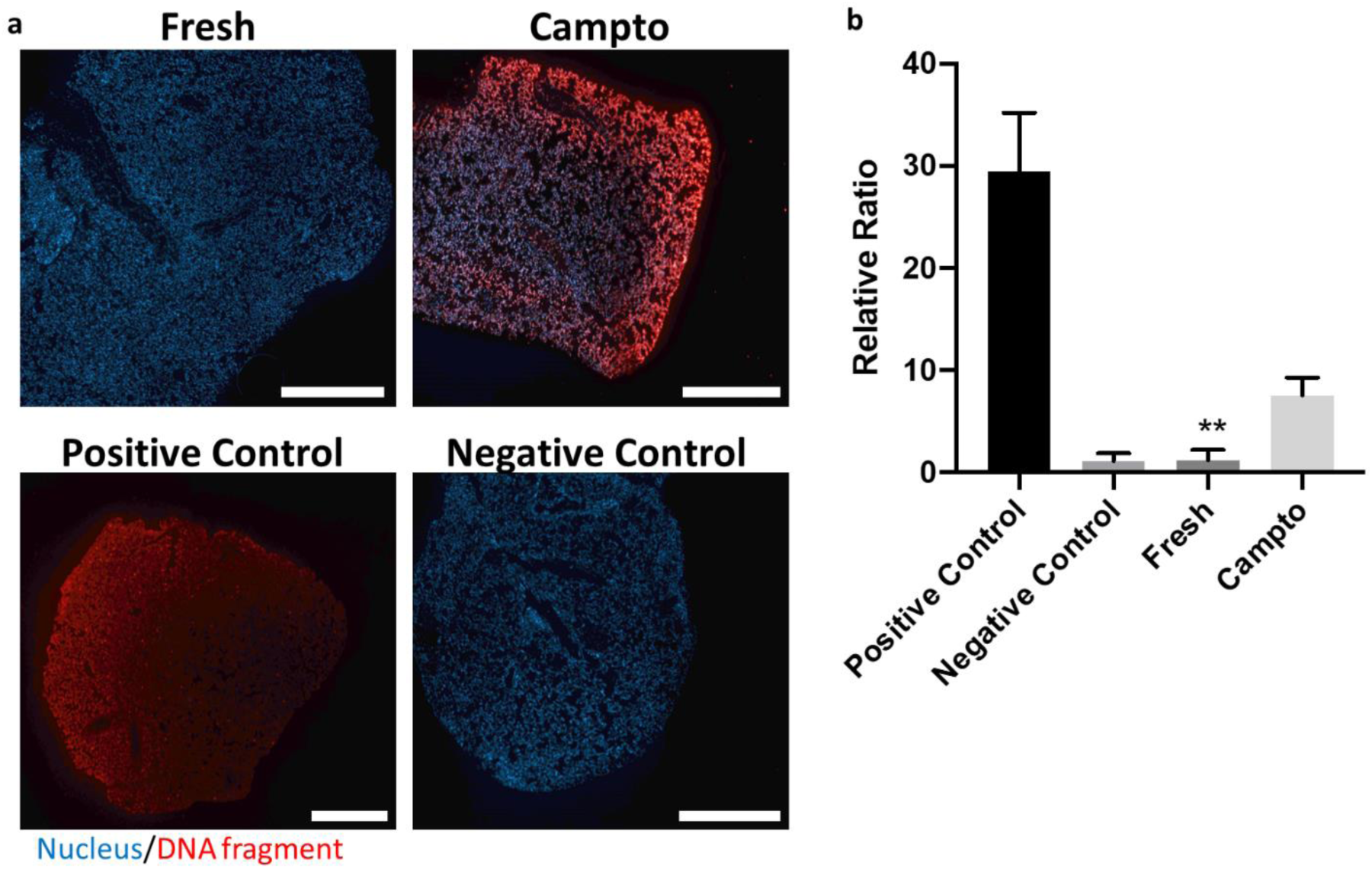
**a**: Confirmation of apoptosis in lung tissue-resident cells via commercially available terminal deoxynucleotidyl transferase dUTP nick end labeling (TUNEL) assay. Blue: DAPI, Red: DNA fragments. Scale bar: 1 mm. **b**: Fluorescence intensity of DNA fragmentation ratio, as defined by the ratio between red and blue. ** p<0.01 vs. Campto.

Camptothecin is an FDA-approved chemotherapy drug that induces apoptosis by inhibiting DNA topoisomerase I^18^. DNA fragmentation is a late-stage event during apoptosis, and can be detected by fluorescence staining of DNA fragments using a TUNEL (Terminal dUTP Nick End-Labeling) assay^19^. This assay was used on camptothecin-assisted decellularization of rat sciatic nerves by Cornelison et al^14^. The authors also reported that apoptosis-assisted decellularized nerves exhibited high cytocompatibility *in vitro* and immunocompatibility *in vivo* after subcutaneous implantation^14^. Another marker of apoptosis is externalization of phosphatidylserine to facilitate phagocytosis of apoptotic cells^19^. Annexin V is another marker of apoptosis as it is bound to phosphatidylserine. This method was used by Bourgine et al. to confirm apoptosis of immortalized human mesenchymal cells in porous ceramic-based 3D scaffolds after decellularization using B/B homodimerizer^15^. To date, these two studies are the only reports of apoptosis-assisted decellularization. These studies collectively demonstrate that apoptosis can be used to decellularize excised tissues and engineered tissue constructs, and warrant further investigation and expansion of apoptosis-assisted decellularization.

### 3.2. Intracellular Components Are Removed and ECM Proteins Are Preserved After Apoptosis-assisted Lung Decellularization

To further assess the extent of decellularization, paraffin sections of lung pieces were stained to image various intracellular components: CD31 for transmembrane protein (Figure 4a-e), alpha smooth muscle actin (α-SMA, Figure 4f-j) for cytoskeletal protein, and DAPI for cell nuclei (Figure 4k-o). As shown in Figure 4, each of these intracellular components remained within the tissue after the camptothecin step. However, a marked reduction in the presence of these components was observed after the SB-10 step, and a significant removal of these intracellular components was observed at the end of decellularization.

**Figure 4.**
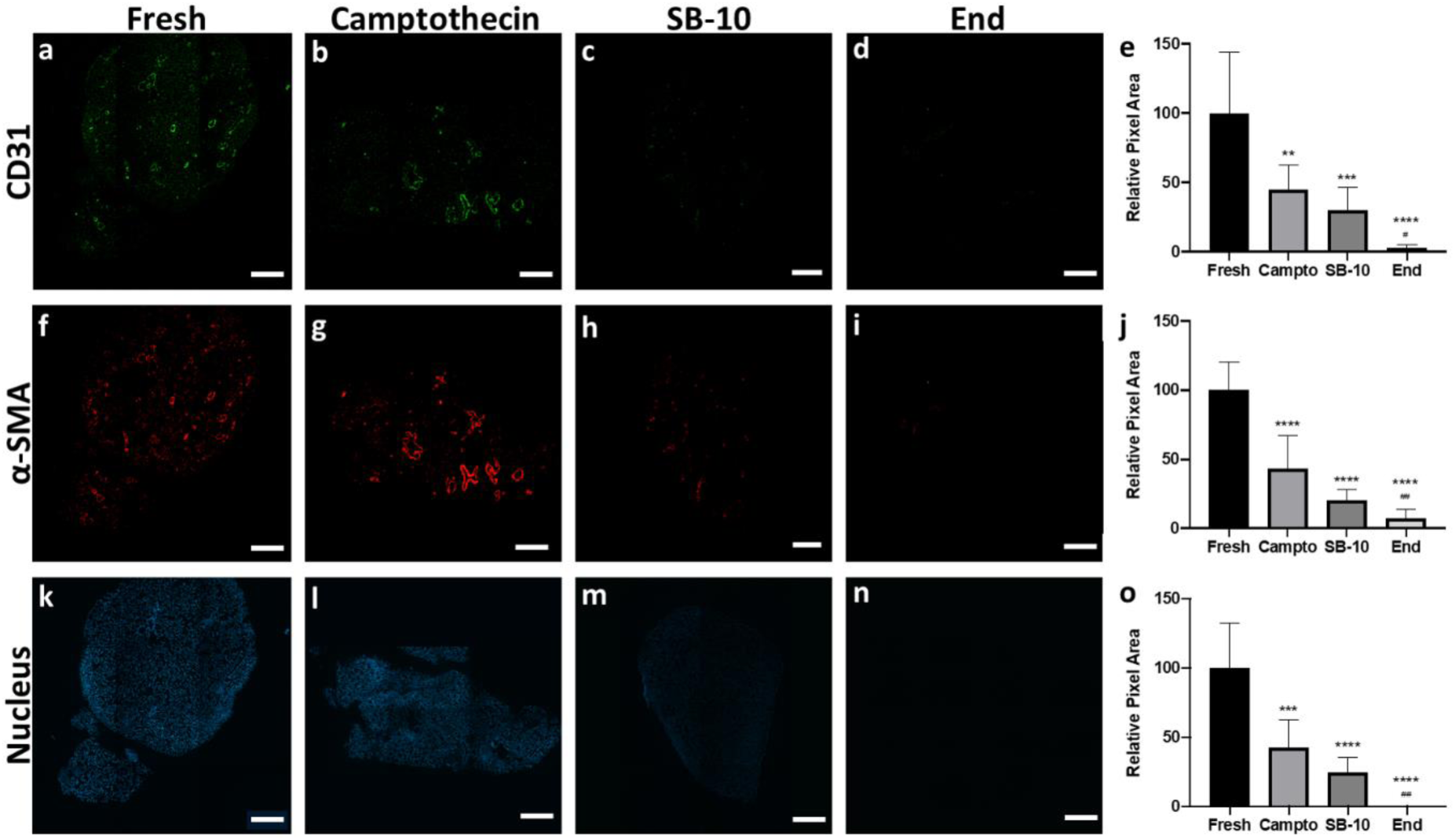
Immunofluorescence analysis of intracellular component removal during the decellularization process. **a, f, k**: fresh lung tissues; **b, g, l**: right after camptothecin treatment, **c, h, m**: right after SB-10 wash, and **d, l, n**: end of decellularization. **e, j, and o** show semi-quantitative analysis of staining intensity for each marker. **a-e**: CD31, **f-j**: alpha-smooth muscle actin, **k-o**: cell nucleus (DAPI). Scale bar: 500 µm. *: p<0.05 vs. Fresh, and #: p<0.05 vs Camptothecin. For each symbol, x indicates p<0.05, xx p<0.01, xxx p<0.005, and xxxx p<0.001.

A key criterion of successful decellularization is not only the removal of intracellular components, but the preservation of extracellular matrix proteins to maintain bioactive cues for both transplantation and *in vitro* test bed development. To this end, the presence of different extracellular matrix proteins was assessed via immunohistochemistry. As shown in Figure 5, collagen I, collagen IV and laminin are all preserved throughout decellularization, indicating that our apoptosis-assisted lung piece decellularization method is not detrimental to preservation of tissue architecture and ECM components.

**Figure 5.**
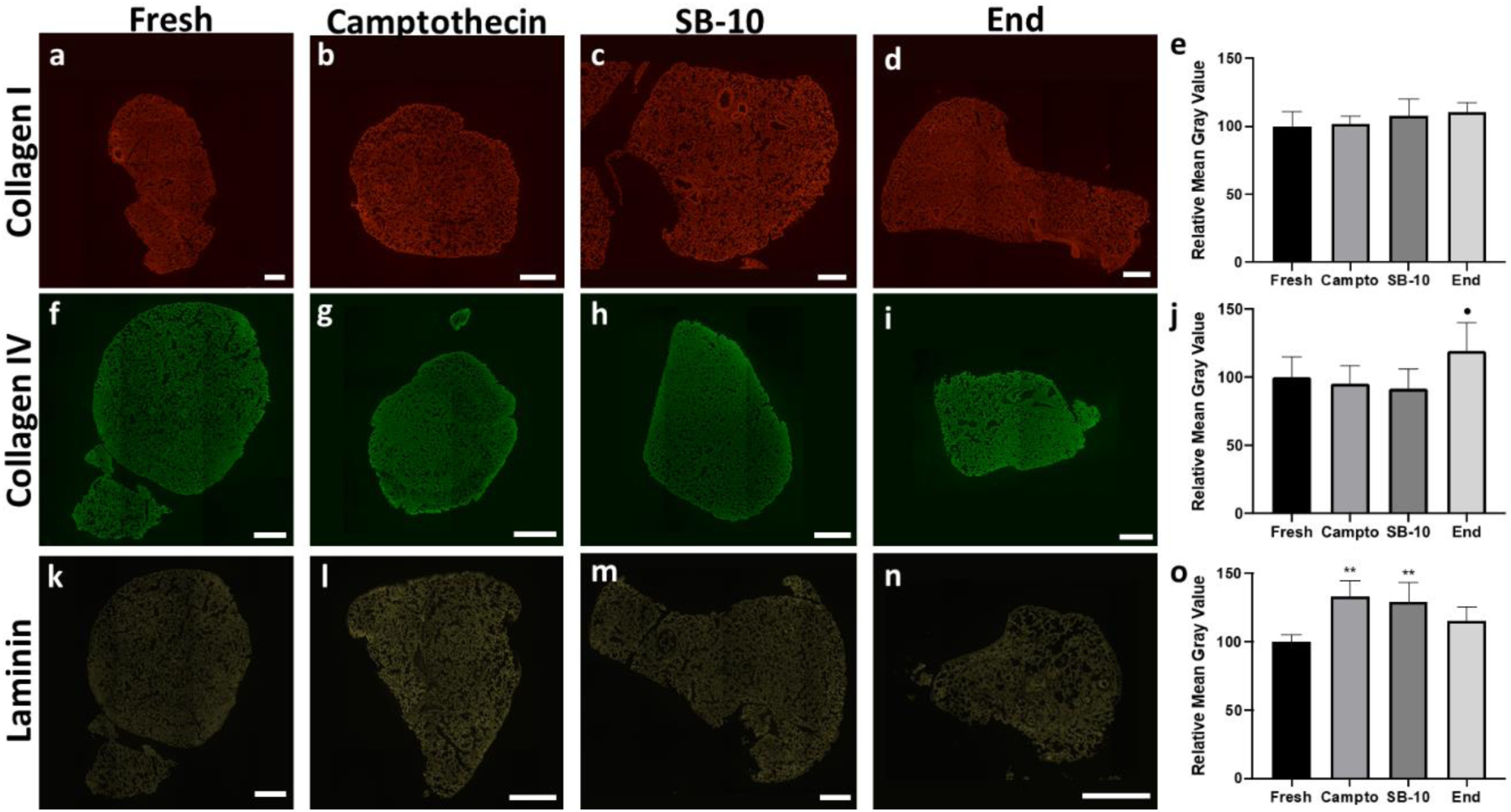
Immunofluorescence analysis of extracellular matrix preservation during the decellularization process. **a, f, k**: fresh lung tissues; **b, g, l**: right after camptothecin treatment, **c, h, m**: right after SB-10 wash, and **d, l, n**: end of decellularization. **e, j, and o** show semi-quantatitative analysis of staining intensity for each marker. **a-e**: collagen I, **f-j**: collagen IV, **k-o**:laminin. Scale bar: 500 µm. *: p<0.05 vs. Fresh, and •: p<0.05 vs Camptothecin. For each symbol, x indicates p<0.05, and xx p<0.01.

Next, the need to use both camptothecin and SB-10 wash verified by comparing three different methods: 1) current apoptosis-assisted method (termed “Campto+SB10”), 2) Campto+SB10 minus the camptothecin treatment (“SB-10 Only”), and 3) Campto+SB10 minus the SB-10 wash (“Campto Only”). As shown in Figure 6, neither Campto Only nor SB-10 Only groups achieved successful removal of all the intracellular components tested compared to the Campto+SB10 method. Specifically, cytoskeletal protein α-SMA removal was significantly better in Campto+SB10 compared to Campto Only, whereas removal of transmembrane protein CD31 and cell nuclei was significantly better in Campto+SB10 compared to both Campto Only and SB-10 Only groups. It is likely that pulmonary surfactants prevented removal of apoptotic bodies without SB-10 washes. Pulmonary surfactants are a complex mixture of lipids and proteins that reduce air-liquid interface surface tension and allows breathing^20^. SB-10 is a zwitterionic detergent that is excellent in disrupting lipid-lipid interactions without significantly damaging the thin protein layer^5^, and hence could have contributed to removing camptothecin-induced apoptotic bodies.

**Figure 6:**
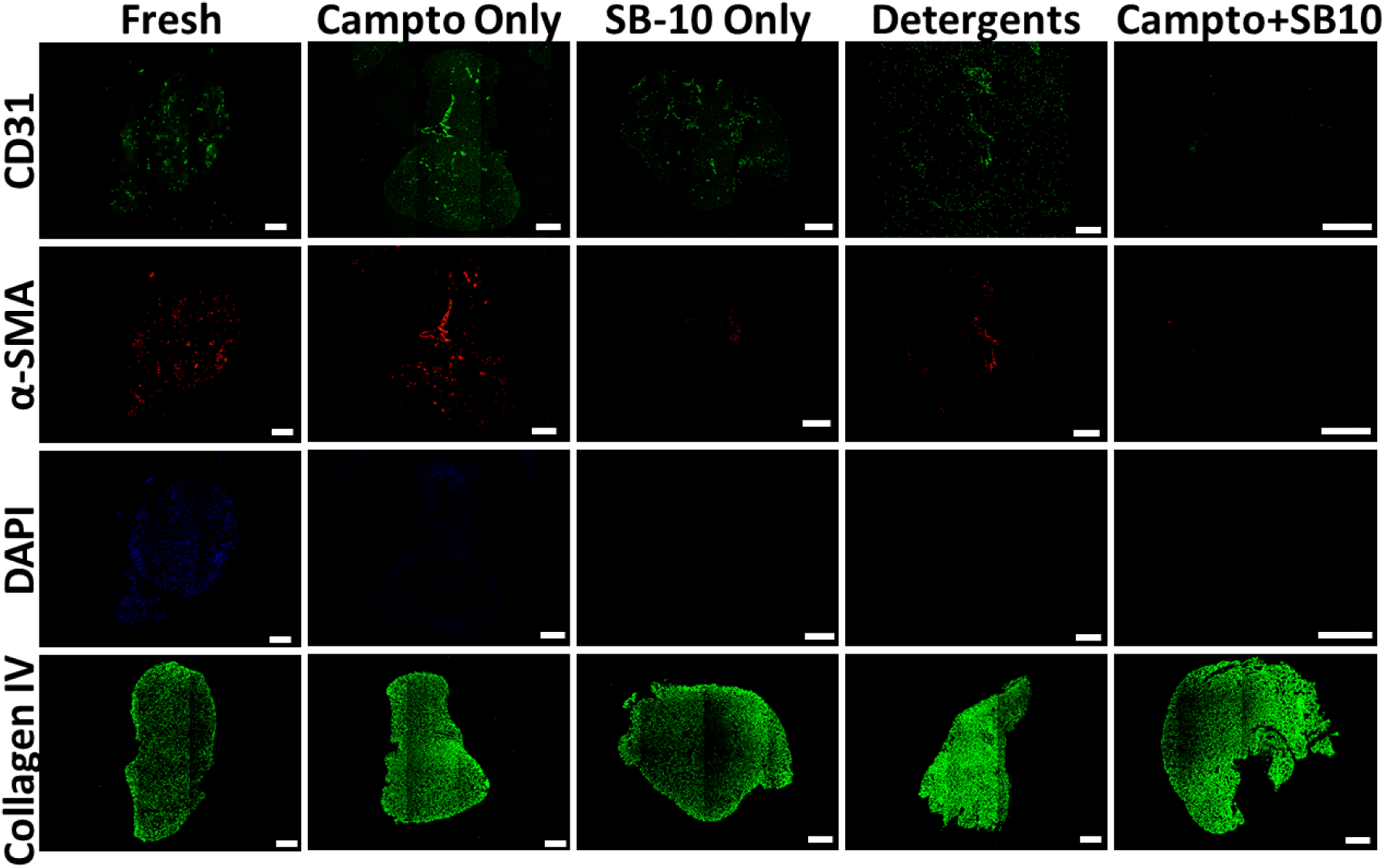
Immunofluorescence images of intracellular component removal and extracellular matrix preservation during different decellularization processes. **Fresh** indicates fresh lung pieces; **Campto Only**: Campto+SB10 minus SB-10 wash. **SB-10 only**: Campto+SB10 minus the initial camptothecin wash. **Detergents**: detergent-based lung tissue decellularization based on a previously established method^13^. **Campto+SB10**: current decellularization method. Scale bar: 500 µm.

Finally, the current method and a previously established, detergent-based decellularization method were compared. There exist numerous reports on detergent-based lung decellularization methods. The most commonly used detergent(s) are 1) sodium dodecyl sulfate and Triton X-100^21–23^, 2) CHAPS^24,25^, and 3) Triton X-100 and sodium deoxycholate^16,26–30^. Among these combinations, we chose Triton X-100 and sodium deoxycholate based on previous reports showing higher cytotoxicity threshold of these detergents compared to others^31^. To this end, we compared the current method to a purely detergent-based method using Triton X-100 and sodium deoxycholate^16^ (referred to as “Detergents”). Compared to the Detergents method, the Campto+SB10 group had significantly less cytoskeletal protein α-SMA as measured by relative percent thresholded area in Figure 7. Additionally, Campto+SB10 and Detergents groups reduced transmembrane protein CD31 levels similarly, while Campto+SB10 had significantly better removal than the SB-10 Only group. Together, these results suggest that both camptothecin and SB-10 together reduce intracellular components (i.e., cytoskeletal and transmembrane proteins) better compared to either SB-10 Only or the Detergents method. The current approach also achieved similar levels of DAPI removal and ECM preservation (measured by average fluorescence intensity of collagen IV) when compared to Detergents (Figure 7). Moreover, Campto+SB10 had significantly higher average fluorescence intensity for collagen IV than SB-10 Only.

**Figure 7.**
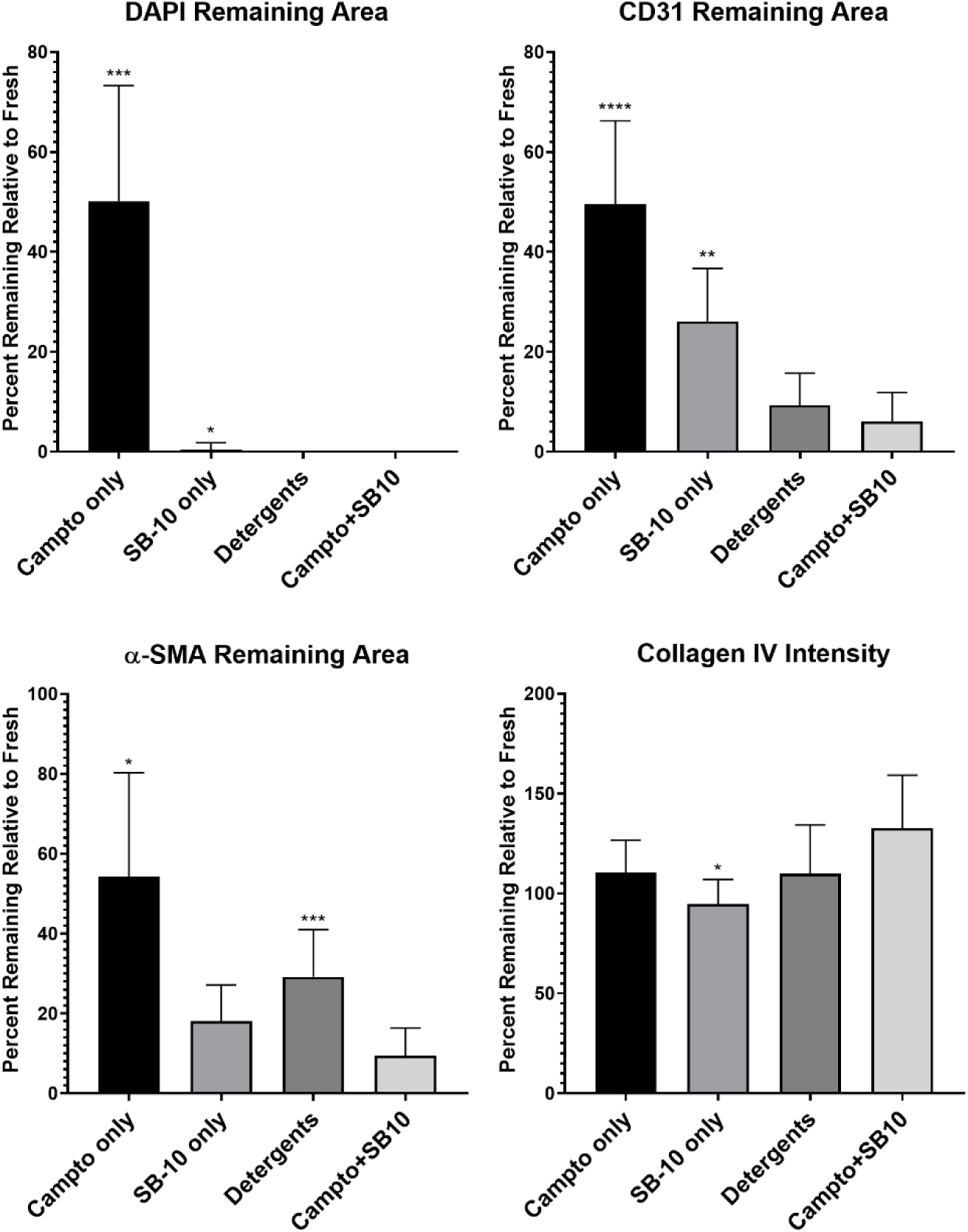
Semi-quantitative analysis of nuclear (DAPI), transmembrane (CD31), and cytoskeletal (α-SMA) percent thresholded area remaining and of extracellular (collagen IV) average fluorescent intensity relative to fresh lung pieces. Only statistical differences for Campto+SB10 are shown, *indicates p<0.05, **p<0.01, ***p<0.001, and **** p<0.0001, n=7-8/group

### 3.3. Quantitative Assessment of Decellularization

To further validate immunohistochemical analysis, quantitative biomolecular assays were performed. DNA, glycosaminoglycan (GAG), and collagen contents were measured from the Fresh, Campto+SB10, SB-10 Only, and Detergents groups. All three decellularization methods significantly decreased DNA content normalized to dry tissue weight (Figure 8a). Importantly, the Campto+SB-10 decellularization method yielded DNA content of less than 50 ng dsDNA/mg dry tissue, which is one of the minimal criteria of satisfactory decellularization based on *in vivo* responses^5^. The lack of visible nuclei from H&E and DAPI staining also meet the criteria, which was observed in the current method in Figures 2 and 4, respectively.

**Figure 8.**
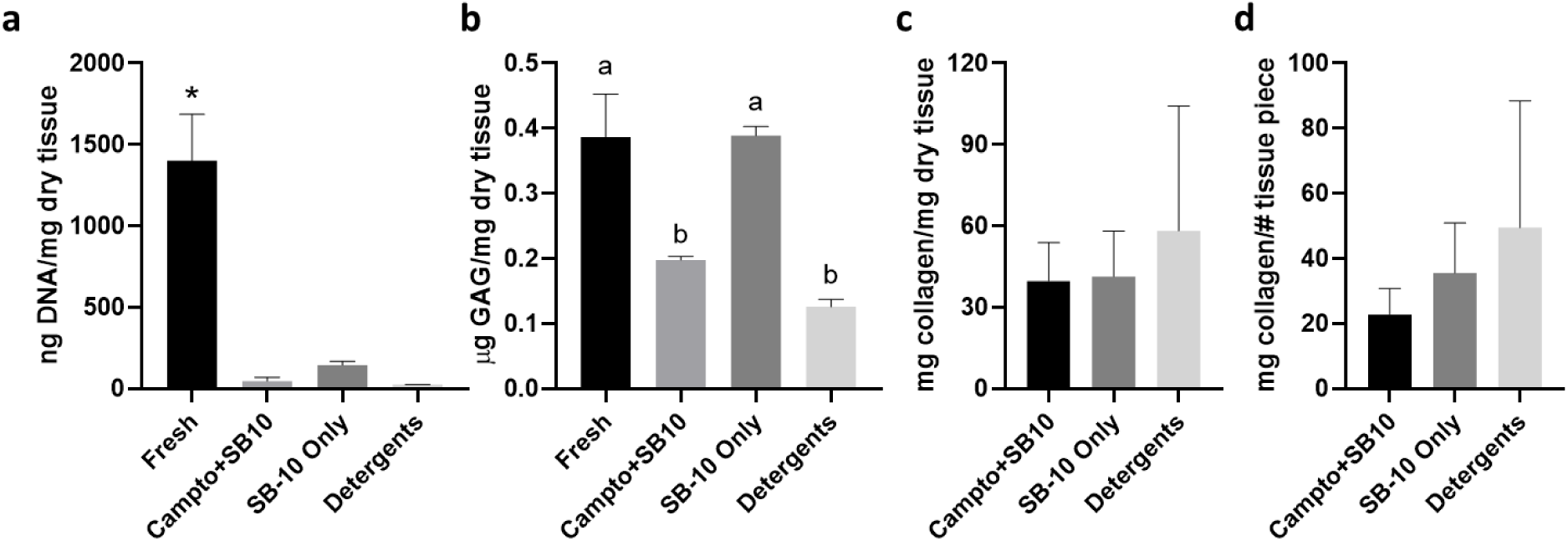
Quantitative assessment of nuclear removal and extracellular matrix preservation after different decellularization methods. In all graphs, Fresh: fresh lung tissues, Campto+SB10: current decellularization method, SB-10 Only: Campto+SB10 minus the initial camptothecin wash, and Detergents: a detergent-based lung decellularization. **a**: DNA content in the decellularized lung tissues, normalized to dry tissue weight in mg. * indicates p<0.05 vs. all other groups. **b**: glycosaminoglycan (GAG) content in the decellularized lung tissues, normalized to dry tissue weight in mg. Different letters indicate statistical significance. **c**: collagen content in the decellularized lung tissues, normalized to dry tissue weight in mg. **d**: collagen content in the decellularized lung tissues, normalized to the number of the decellularized lung tissues used per group.

Interestingly, Fresh and SB-10 Only groups had similar and significantly higher GAG content than the Campto+SB10 and Detergents groups (Figure 8b). A decrease in GAG content after decellularization has been reported in other studies. Wagner et al. decellularized healthy and emphysematous human lungs using 0.1% Triton X-100 and 2% sodium deoxycholate and observed GAG reduction regardless of healthy versus diseased states of the excised lungs^32^. Consistent with this study, Balestrini et al. also saw a significant decrease in GAG content in porcine lungs decellularized with the same concentrations of both of the detergents; however, lower concentrations of both Triton X-100 (0.0035%) and sodium deoxycholate (0.01%) preserved GAGs^30^. GAG decrease after decellularization is observed in other tissues, including liver (sodium dodecyl sulfate)^33^, pericardium (Triton X-100)^34^, pulmonary valve leaflets (trypsin-EDTA)^35^, and fibroblast culture sheets (sodium dodecyl sulfate)^36^. Nevertheless, these decellularized matrices were successfully recellularized and remained functional.

Furthermore, normalized collagen content was not statistically different in all decellularization methods (Figure 8c). Preservation of collagen is key, because it is not only the most abundant ECM protein in human tissue, but also thermally-gelation of the hydrogel is majorly dependent on self-assembly of collagen I fibrils^37^. In this study, hydroxyproline content was measured as a means of quantifying collagen content. Hydroxyproline is a nonessential amino acid that makes up approximately 13.5% collagen^38^. Given the high hydroxyproline content in collagen, hydroxyproline measurement has been used in determining collagen content in a variety of tissues including lungs^39–42^. Furthermore, urinary hydroxyproline is used as a main marker of osteoporosis, indicating collagen degradation from bone resorption^43^. Since tissue decellularization leads to a significant reduction in tissue weight^44^, as shown in Figure 9a, collagen content was also normalized against the number of tissue pieces. As shown in Figure 8d, the collagen content was not statistically different across different groups. To determine what ECM proteins are present in the tissues after decellularization, proteomic analysis can be performed^45^, e.g., mass spectrometry and Western blots.

**Figure 9.**
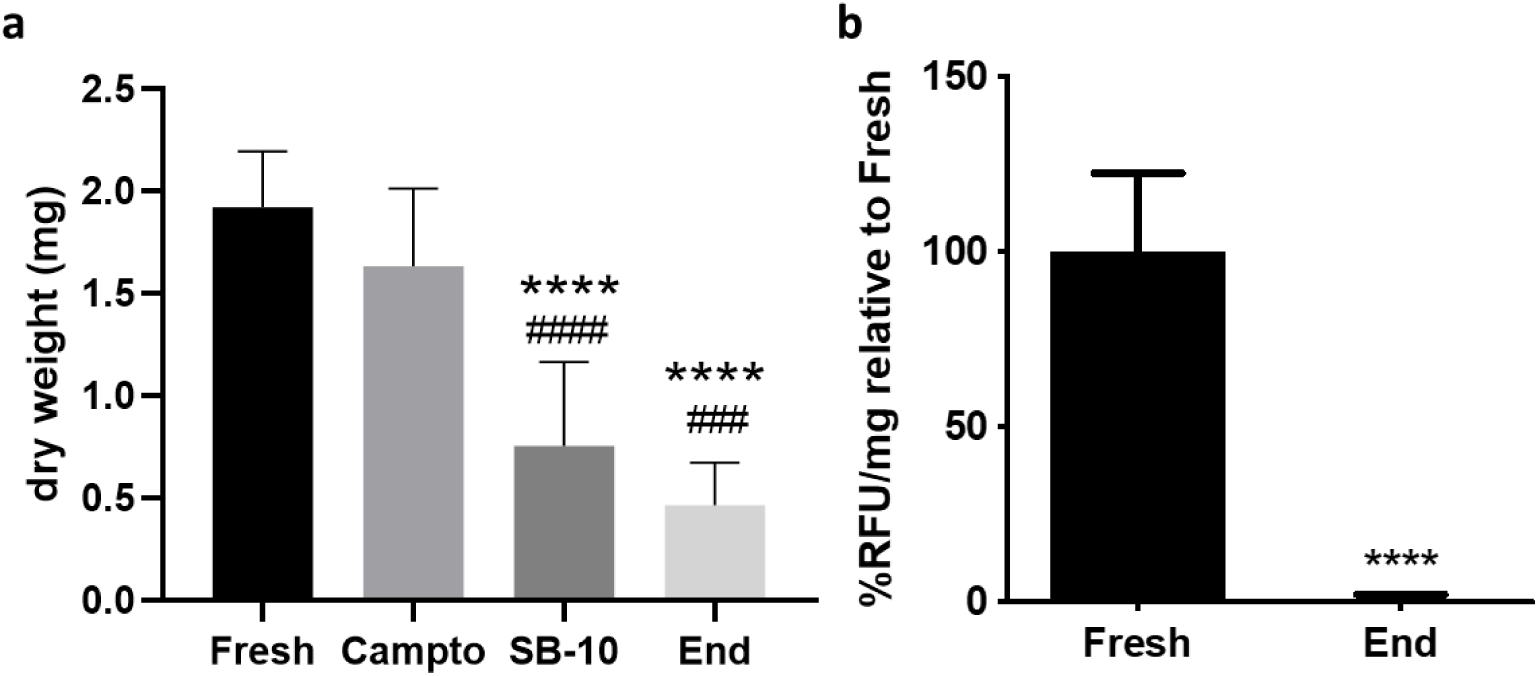
**a**: dry tissue weight at the different stage of the current decellularization method. * p<0.05 vs. Fresh, # p<0.05 vs. Campto. **b**: fluorescence intensity of the media incubated with fresh and decellularized lung pieces, as an indicator of metabolic activity in the resident cells using alamarBlue assay. **** p<0.001.

Finally, tissue weights at different steps of the Campto+SB10 method were measured as an indicator of cell removal. As discussed earlier, a significant reduction in tissue weight was obtained at the end of decellularization, corresponding with the changes in DNA, GAG and collagen contents. Finally, to verify that there are no viable, residual cells after decellularization, metabolic activity of the lung pieces was measured using alamarBlue assay. As shown in Figure 9b, decellularized lung pieces showed no signs of metabolic activity, indicating successful removal of viable, functioning cells. The alamarBlue assay reagent contains a low fluorescence molecule resazurin, which is converted by metabolically active cells into resorufin, a high fluorescence molecule. The shift in fluorescence of the culture media is then detected and correlated with metabolic activity of the cells. In this study, fresh lung pieces contained metabolically active cells, which were undetected after the current decellularization method.

### 3.4. Lung ECM Hydrogel Formation and Rheological Property Characterization

To create biocompatible 3D hydrogels, decellularized lung pieces were lyophilized and digested in pepsin-acid solution. Afterwards, pH level and osmotic balance of the pre-gel solution were matched to physiological values and incubated at 37°C (Figure 10a). As shown in Figure 10b, translucent lung ECM hydrogel was formed. Further, the lung ECM hydrogel had linear viscoelastic characteristics in the plateau region between 0.1 and 3.98 rad/s. Within the linear viscoelastic range, the lung ECM hydrogel demonstrated a larger storage modulus (G’) than loss modulus (G′′) at 37°C (Figure 10c). This relationship indicates that our lung ECM hydrogel has a solid-like behavior throughout the range of frequencies, as observed in other previously reported ECM-based hydrogels^46,47^. Overall, these results indicate that our lung ECM hydrogels are uniform and stable, thus suitable for 3D cell culture^48^.

**Figure 10.**
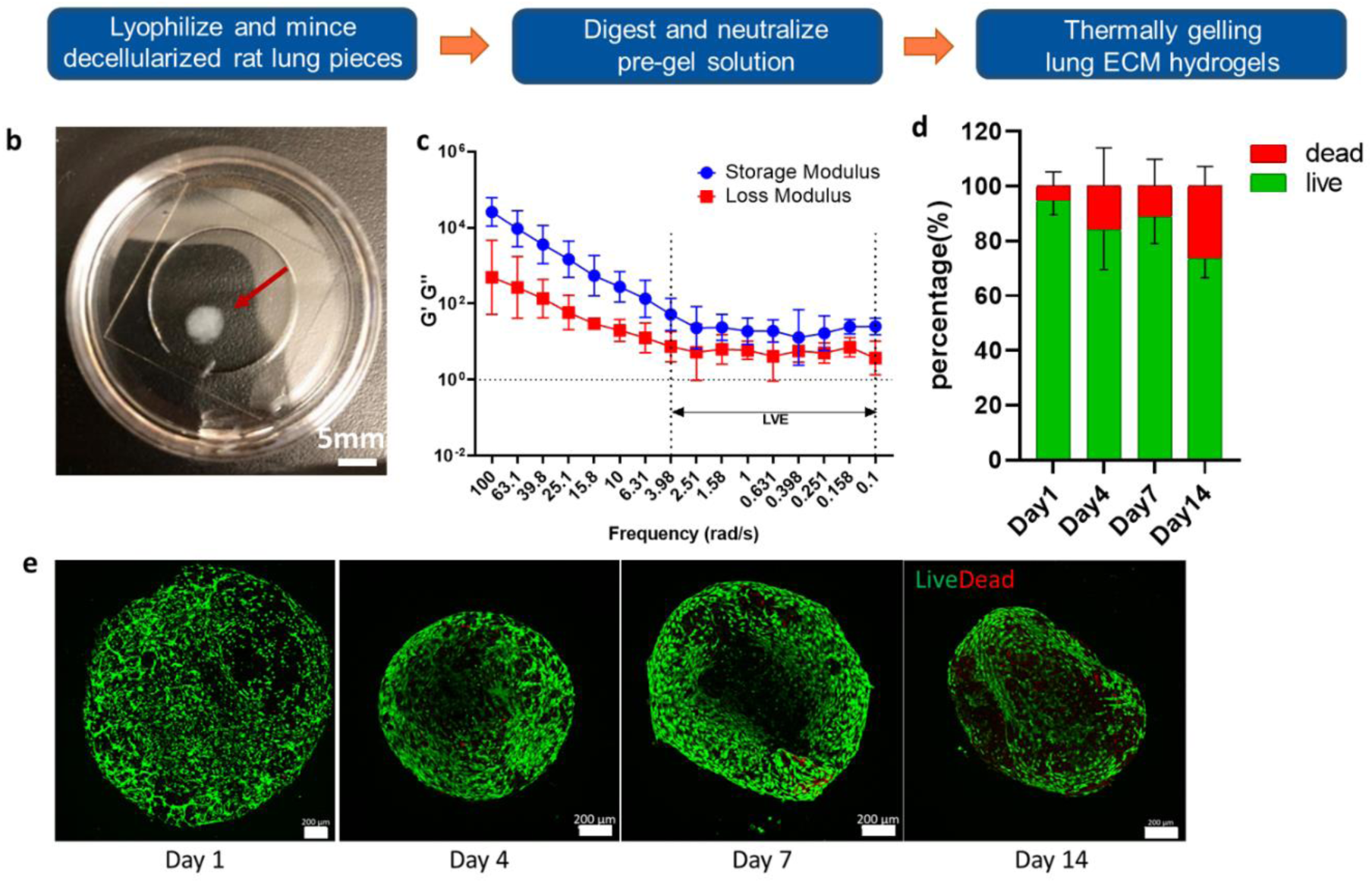
**a**: overview of thermally-gelling lung ECM hydrogel fabrication process. **b**: an example of the lung ECM hydrogel. **c**: Viscoelastic properties of the lung ECM hydrogel as measured by frequency sweep. **d**: quantification of live vs. dead CCL-149 rat pulmonary epithelial cells in the lung ECM hydrogel, as shown in panel e. **e**: representative confocal images of CCL-149 embedded in 3D lung ECM hydrogel. Images taken at days 1, 4, 7 and 14. Scale bar: 200 µm.

### 3.5. Pulmonary Epithelial Cell Viability in Lung ECM Hydrogel in vitro

To confirm that high cell viability can be maintained inside the lung ECM hydrogels, CCL-149 rat pulmonary epithelial cells were cultured inside the hydrogels, and viability was measured over two weeks using a live/dead assay. As presented in Figures 10d-e, these hydrogels were biocompatible as they supported long-term 3D culture of CCL-149s with high viability maintained throughout the culture periods.

These lung ECM hydrogels are versatile scaffolds that can be used for disease modeling. Several studies have reported the feasibility of decellularized lung ECM hydrogels for studying metastatic behavior of lung cancer cells^22,49,50^. Furthermore, decellularizing healthy versus diseased lungs also altered seeded cell behavior, including increased fibrotic marker expression^51,52^, epithelial-to-mesenchymal transition of pulmonary epithelial cells^53^, and poor cell adhesion^32^.

High cell viability in lung ECM hydrogels indicates the possibility of use of lung matrices for regenerative medicine. A study by Wu et al. delivered nebulized pre-gel lung ECM solution via inhalation and observed that it rescued hyperoxia-mediated damage, including apoptosis and oxidative damage of pulmonary cells^54^. Similarly, a study by Link et al. generated lung ECM nanoparticles via electrospraying of the pre-gel solution and observed that resulting lung ECM nanoparticle promoted pro-regenerative M2 polarization of macrophages *in vitro*^29^.

Our current study demonstrated that apoptosis-assisted decellularization can be achieved in rat lung tissues. The immediate next step is to scale up the decellularization to whole lungs. Most lung decellularization methods utilize perfusion of detergents and washing buffers^16,21,23,25^; however, a study by Nagao et al. revealed that perfusion led to increased leakiness from pulmonary capillary beds compared to a diffusion method, in which lung lobes were simply soaked in solutions with agitation^17^. A thorough examination of tissue integrity with different delivery of decellularization solutions will need to be performed. Once completed, apoptosis-assisted decellularized lungs can be implanted *in vivo* subcutaneously for immune response^14^ as well as orthotopic transplantation to replace damaged lungs. Recellularization of decellularized lungs will also need to be assessed to create fully functioning transplantable lungs^21^.

Furthermore, the lung ECM hydrogels can be further utilized to create *in vitro* test beds of different pulmonary pathological conditions, including lung cancer, asthma, chronic obstructive pulmonary disease and pulmonary cystic fibrosis. In particular, microfluidic models of healthy and diseased lungs can incorporate our hydrogels to study gas exchange between vascular and alveolar beds and other pulmonary (patho)physiological events in a physiologically relevant manner^55,56^.

## 4. Conclusion

This study presents a novel method of lung tissue decellularization by inducing apoptosis using camptothecin and washing the cellular debris using SB-10. We show that our method results in comparable preservation of ECM components and better removal of intracellular components than a currently existing detergent-only based method. Furthermore, these decellularized lung tissues can be further processed into 3D hydrogels (injectable, in situ gelling) where encapsulated cells maintain high viability over two weeks. This novel method of tissue decellularization can be extended to other tissues and organs and avoid the heavy use of detergents to remove cellular components. Future work includes scaling up to whole organ levels, assessing *in vivo* response to tissues and organs decellularized via apoptosis, and creating *in vitro* disease models to improve our understanding of pathological conditions in a native-like platform.

## Acknowledgements

This work was funded by the National Science Foundation through Award Number 1605223. We would like to thank Dr. Edward Phelps and his lab members at the University of Florida for sharing their pressure cooker for the antigen retrieval step in immunohistochemistry. We would also like to thank Dr. Benjamin Spearman for helpful discussions on rheology.

